# EXTRANEURAL RABIES VIRUS INFECTION LEADS TO TISSUE DAMAGE AND CELL DEATH IN MICE

**DOI:** 10.1101/2023.05.04.539354

**Authors:** Érika D. Leal Rodrigues, Vinicius Pacheco da Silva, Victor G. Bastos Chaves, Cássia N. de Sousa Moraes, Sara de Souza Pereira, André L. Nogueira Lima, Taciana Fernandes Barbosa Coelho, Pedro F. da Costa Vasconcelos, Ana C. Ribeiro Cruz, Livia Medeiros Neves Casseb

## Abstract

Rabies, a fatal neurological disease caused by *Lyssavirus rabies* (RABV), poses a significant threat to public health globally. Despite extensive studies on RABV-induced neuropathology, the involvement of extraneural organs during rabies pathogenesis and the tropisms of wild-type strains to different organs remain largely unknown. Here, we investigated the tropism of a dog and bat RABV variant to three different extraneural tissues (kidneys, lungs and liver) and characterized cellular and tissue damage associated with infection in mice over 30 days. Our results reveal that RABV may have a tropism for the kidneys and cause tissue-specific cellular damage. Furthermore, we propose that RABV spreads to extraneural tissues simultaneously with central nervous system (CNS) infection. Understanding the involvement of extraneural organs in rabies pathogenesis may contribute to the development of effective treatment strategies of this fatal disease.

**AUTHOR SUMMARY:** Rabies is a lethal viral infection that targets the nervous system and generally can be transmitted to humans by bites of infected animals. While there has been significant research focused on how the virus damages the brain, little is known about how the infection affects other organs in the periphery. To address this knowledge gap, we conducted an experimental study to investigate the effects of two distinct wild strains of the virus, one isolated from dogs and the other from bats, on the lungs, liver, and kidneys in mice model of infection. Our findings suggest that the rabies virus infection leads to cell death and produces specific lesions in each of these organs, and we hypothesize that rabies virus may spread to these tissues at the same time as the brain, which possible contributes to the disease outcome. These findings enhance our understanding on how rabies virus targets organs outside the nervous system and its pathology in these different systems.

## INTRODUCTION

Rabies virus (RABV) is a highly lethal neurotrophic agent that causes severe encephalomyelitis in mammals (1). In the central nervous system (CNS), RABV triggers pro-apoptotic pathways in immune cells to subvert host immune response (2,3) while using anti-poptotic strategies to protect host cells (4). This subversion creates a paradox in which, despite severe neural dysfunction, histopathological brain alterations are relatively mild in patients with the disease (5).

Overall, viral transmission generally occurs through bites from an infected animal to other mammals, including humans (6). Within the host, RABV crosses the neuromuscular junction and binds to several neuronal cellular receptors (7). Hence, RABV migrates centripetally to the CNS through the nerves of the peripheral nervous system (PNS) (8). On the other hand, RABV may also migrate from the CNS to peripheral organs through the same pathway (9). Whereas the viral cycle in rabies is commonly associated with neuroinvasion and CNS infection, RABV can spread to tissues outside the neuronal environment (10).

Several studies have detected RABV in the cornea (11), lungs (12), kidney (13), liver (14), salivary gland (15), and hair follicles (16). Although rarely, the main form of human-to-human viral transmission occurs via organ and tissue transplantation from RABV-infected donors (17), thus suggesting the permissiveness of different cellular populations to the virus Despite the detection of RABV antigen and RNA in many extraneural tissues, the precise kinetics and effects of the infection into these tissues remain poorly understood. It is unclear whether viral load or different RABV strains are relevant to cause tissue damage into peripheral organs. Therefore, our study aimed to characterize rabies-induced tissue damage in extraneural organs and assess whether these damages differ according to the organ. We have quantitatively assessed RABV antigen, morphological changes and cellular alterations in the kidney, lungs, and liver of rabies-infected mice during disease onset, by comparing the effects of two wild-type RABV variants. Our results suggest the commitment of extraneural organs in a tissue-specific manner and provide new insights into RABV tropism and the potential impact of non-neural infection on the disease outcome.

## RESULTS

### Rabies virus kinetics and the cellular tropism in extraneural tissues

We assessed the cellular expression of the viral antigen in target cells of the lung, liver, and kidney throughout 28 days post-infection (d.p.i) to understand the dynamics and tropism of these two different RABV strains in extraneural tissues (Fig 1). Upon the muscular infection, we identified the local expression of viral markers from the 5th d.p.i, which remained expressed in the lung, kidney, and liver until the last day of analysis (Fig 2A).

**Fig 1.**
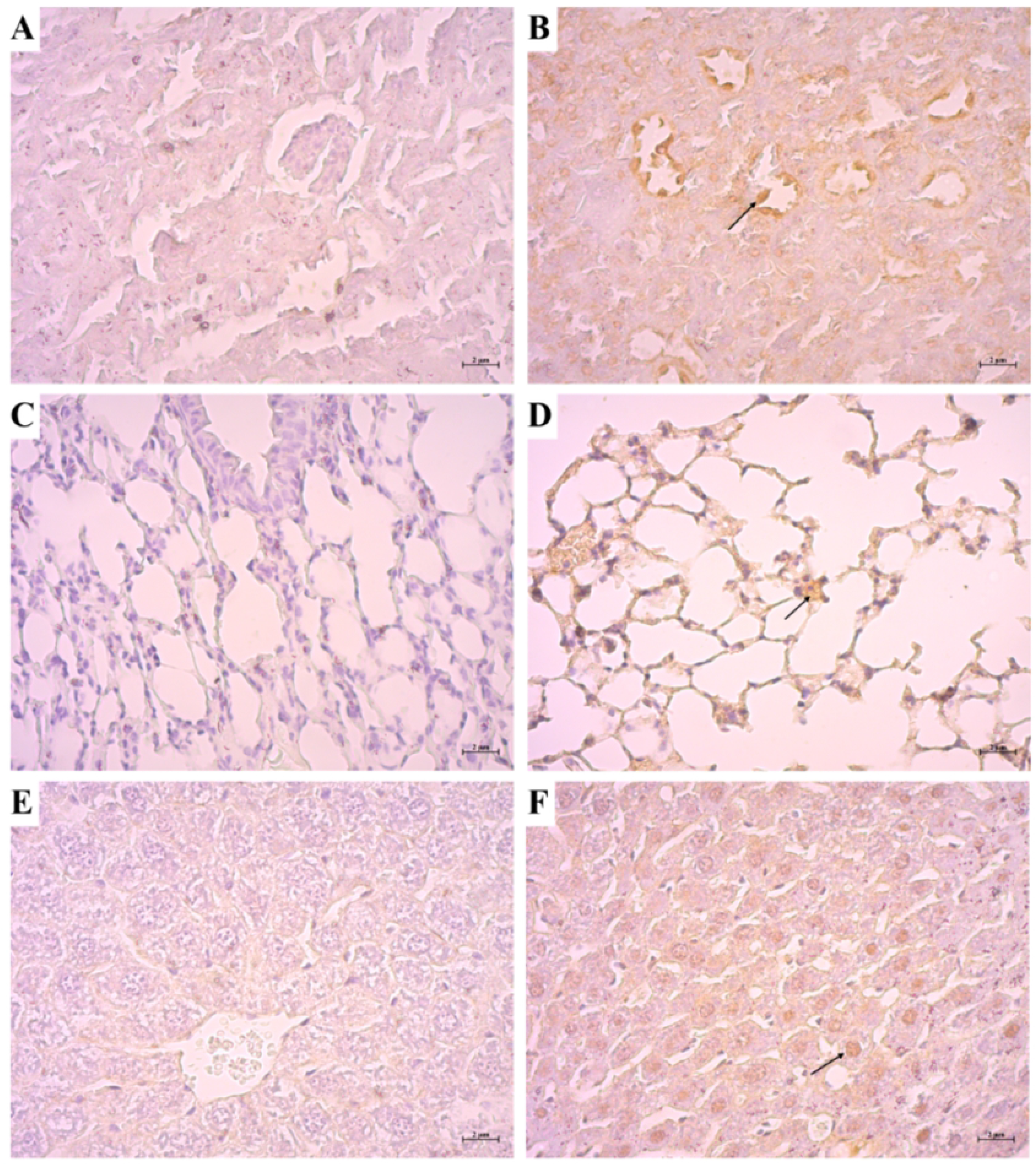
Immunohistochemistry of rabies virus (RABV) antigen in extraneural tissues (brown-colored cells). A) Negative control in the kidney. B) RABV-positive cells in the kidney. C) Negative control in the lung. D) RABV-positive cells in the lung. E) Negative control in the liver. F) RABV-positive cells in the liver.

**Fig 2.**
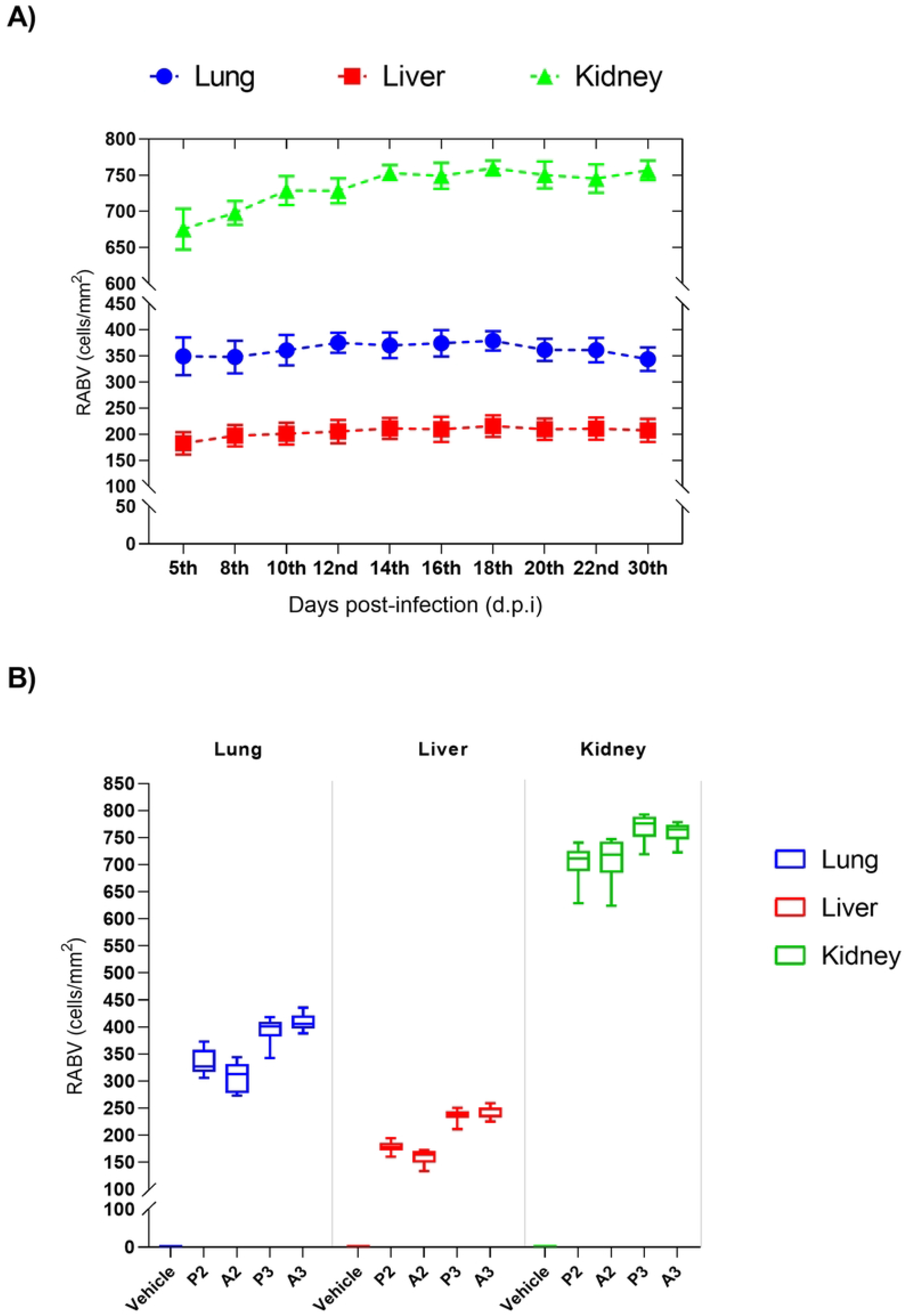
Spatiotemporal rabies virus kinetics in the extraneural tissues. A) Temporal expression and comparative analysis of RABV antigen in the lung (blue), liver (red), and kidney (green). *\*\*\*p ≤ 0*.*001 (kidney vs liver)*; B) RABV differential expression in the lung, liver, and kidney.

When investigating RABV cellular tropism, the kidney was more permissive to the infection in comparison to the liver (Fig 2A) and lung. We also suggest that the viral activity might be affected by RABV variant, since the antigenic-variant-3 (AgV3) – the bat variant – inoculated both anteriorly (A3) and posteriorly (P3), was more expressed than antigenic-variant-2 (AgV2) – the dog variant – also inoculated anteriorly (A2) and posteriorly (P2) – as shown in the Fig 2B.

### Wildtype RABV infection in the kidney induces parenchymal and cellular damage at different degrees

The semiquantitative histopathology analysis (Fig 3) revealed that after the infection, the lesions ranged from moderate to severe, according to a determined score (Fig 4).

**Fig 3.**
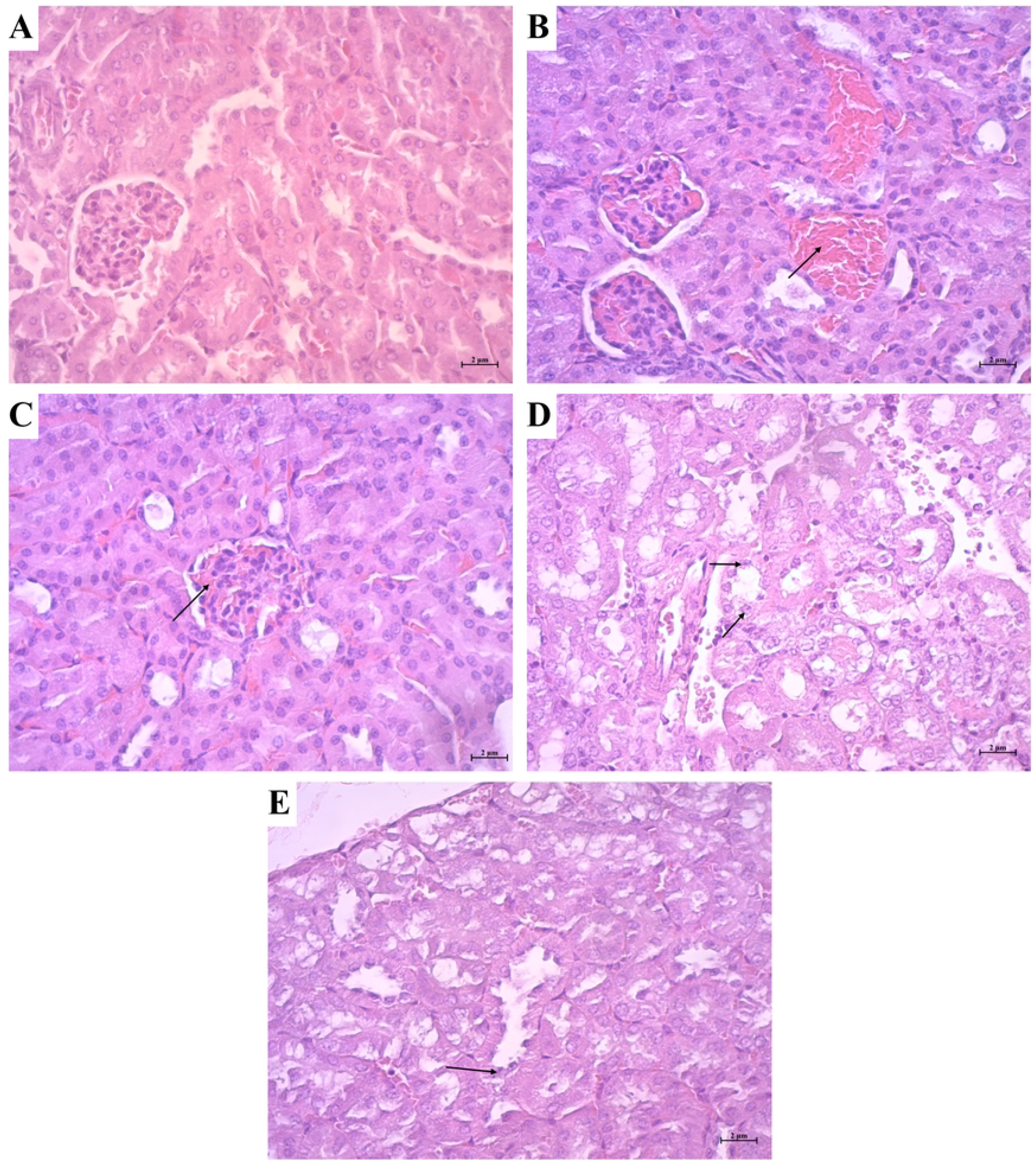
Cellular damage caused by RABV in the kidney (arrows), analyzed by HE staining. A) Kidney negative control. B) Congestion in the kidney parenchyma. C) Glomerular congestion. D) Pyknosis of proximal tubules. E) Pyknosis of distal tubules.

**Fig 4.**
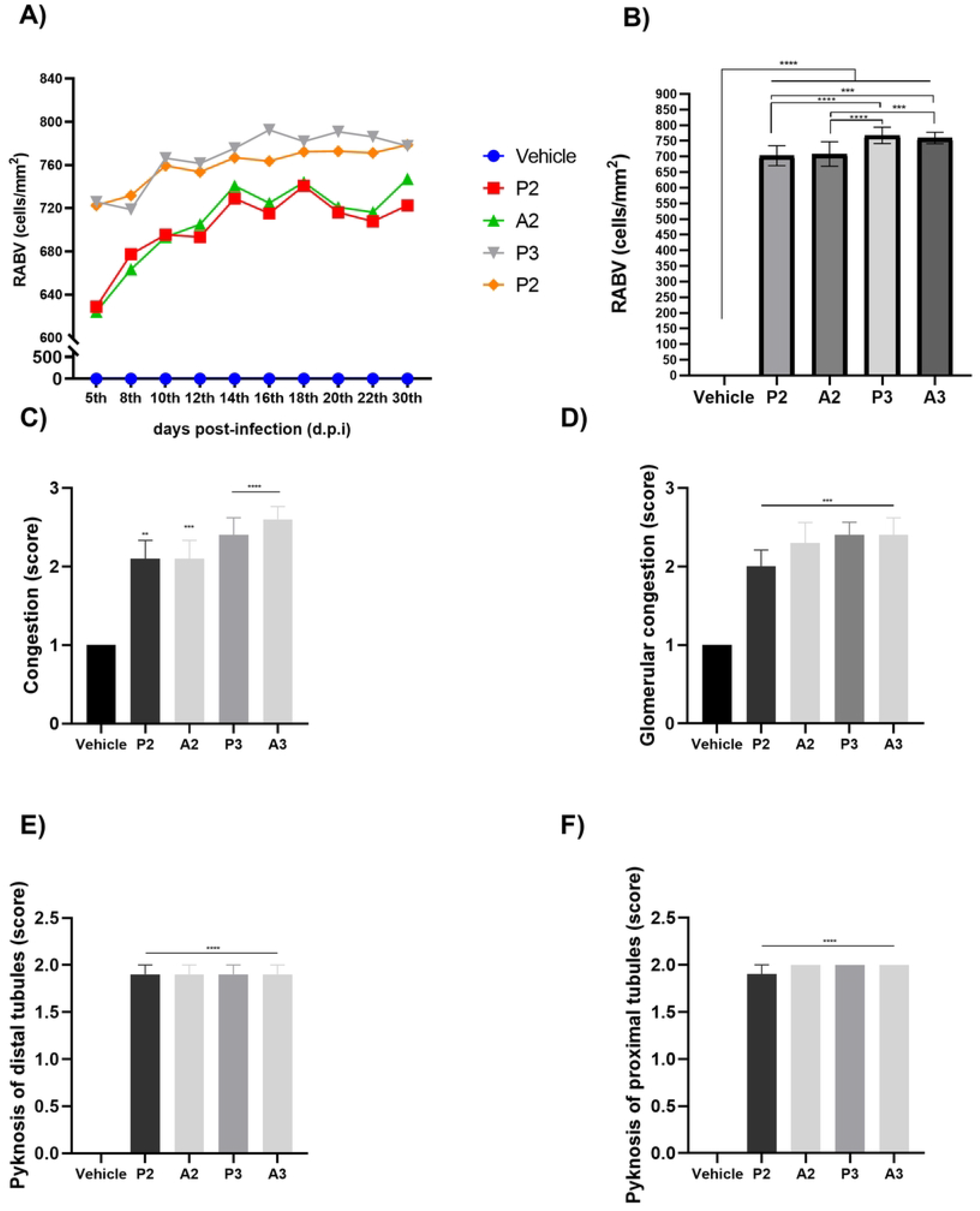
*In situ* rabies expression and RABV-induced damage in the murine kidney. A) Temporal kinetics of RABV-expressing cells; B) Total expression of viral antigen; C) Tissue congestion; D) Glomerular congestion; E) Pyknosis of distal tubules; F) Pyknosis of proximal tubules. \*\*\**p< 0*.*001, ****p<0*.*0001*

The anti-RABV analysis demonstrated that the bat variant was more prominent in the tissue (Fig 4B), and the kinetics indicates a progressive and high viral expression in the cells following the disease progression (mean P2 = 702.7±10.09; A2=708.0±12.33; P3 =767.7±8.18; A3=759.3±5.85 vs vehicle, *p<0*.*0001*) (Fig 4A). Interestingly, the inoculation site may affect the viral activity among these different strains, since the posterior and anterior-infected mice differ among the groups (mean diff. P2 vs P3 = 65.05; A2 vs. A3= 51.31, p<0.01).

When analyzing the parenchymal disorders evoked by the infection with bat and dog variants, we could not observe those differences among the groups, but all infected mice displayed the highest rate of parenchymal and cellular damage compared to the non-infected mice, such as congestion, glomerular congestion, and pyknosis in cells of the distal and proximal tubules of the kidney (Fig 4C-F). Therefore, our data suggest that although AgV3 was more prevalent in the kidney, the viral activity was not necessarily accompanied by the severity of the lesions.

### Bat and canine RABV infect lung cells and promote *in situ* inflammation

Our histopathological analysis in the lungs (Fig 5) suggests the infection with both strains caused *in situ* inflammation followed by the infiltration of immune cells and tissue congestion that ranged from mild to moderate (Fig 6D-F). It was also associated with mild cellular damage, affecting the alveolar wall (Fig 6C).

**Fig 5.**
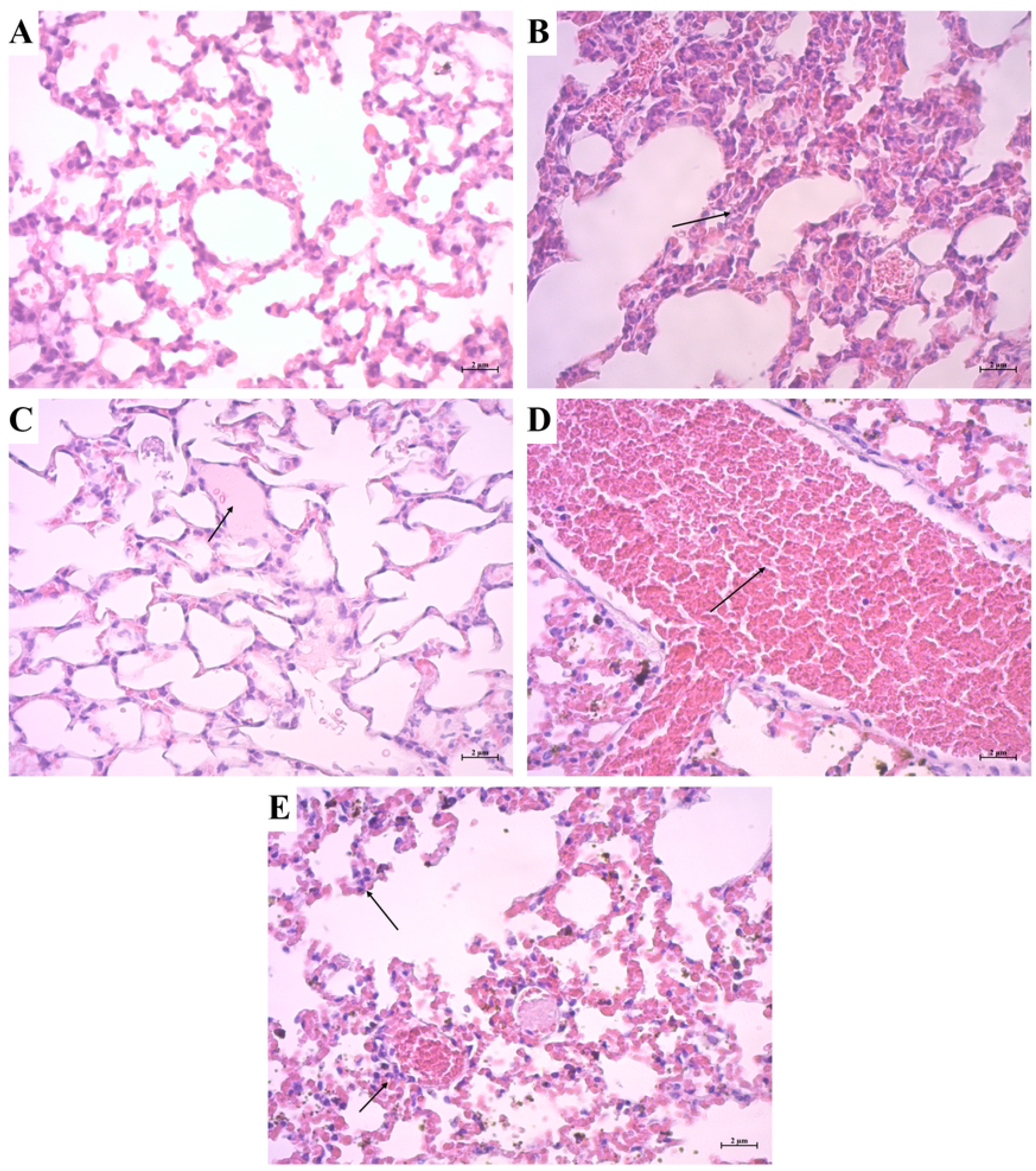
Cellular damage caused by RABV in the lungs (arrows), analyzed by HE staining. A) Lung negative control. B) Alveolar-wall thickening. C) Intra-alveolar edema. D) Major congestion. E) Infiltrative cells in the lung parenchyma.

**Fig 6.**
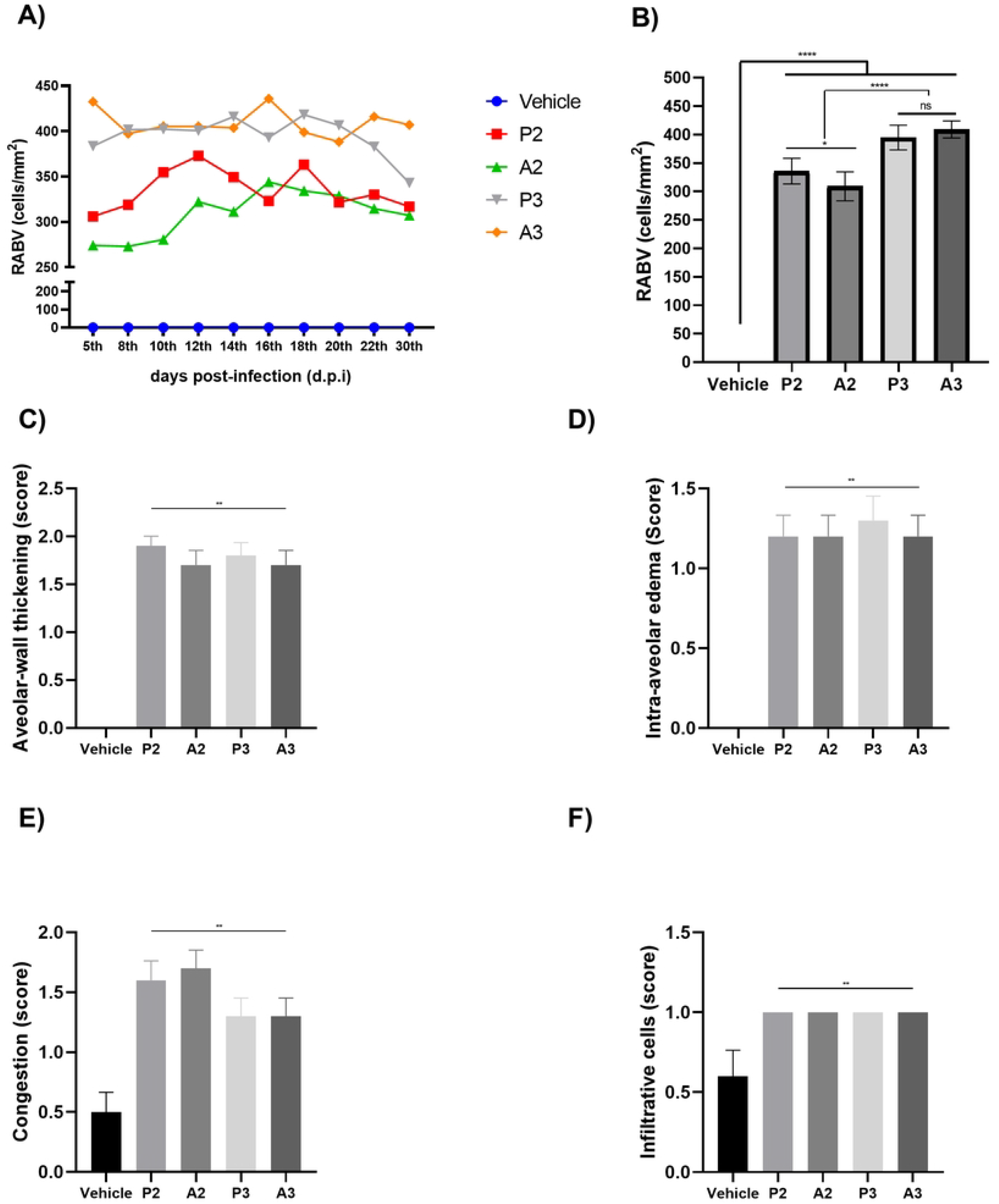
*In situ* rabies expression and RABV-induced damage in the murine lung. A) Temporal kinetics of RABV-expressing cells; B) Total expression of the viral antigen; C) Alveolar wall-thickening; D) Intra-alveolar edema; E) Congestion; F) Infiltrative cells in the parenchyma. \*\**p ≤ 0*.*001*

In the lungs of the experimental groups, we found a similar and progressive expression of both bat and dog-RABV variants from the 5th d.p.i which indicated that, in both cases, the cells that compose the lung parenchyma are permissive to the infection in a lesser extent (mean P2 = 335.7±7.13; A2 = 309.1±8.03; P3= 394.7±6.85; A3 = 409.0±4.76 vs vehicle, *p<0*.*0001*) (Fig 6A). Moreover, unlike the kidney and regarding the bat-variant, the inoculation site may not influence the viral invasiveness since P3 and A3 mice did not differ – but the AgV2 infected mice displayed a different pattern in which the infection site changed the total expression (P2 vs. A2 = 26.62, *p=0*.*0339*) (Fig 6B). It is noteworthy that lung cells appear to be more permissive to infection with the bat-RABV compared to the canine variant.

### RABV infection of Kupffer’s cells enhances cellular damage and pyknosis in the liver without triggering inflammation

The histopathological damages in the liver (Fig 7) varied from mild to severe. Although the liver was infected to a lesser extent, the IHC analysis revealed the expression of RABV antigen from the 5th d.p.i. (Fig 8A), which varied from 120 to 240 RABV-positive cells/mm^2^. Similarly, the expression of AgV3 (P3= 237.3±3.31; A3 = 244.2±3.36 vs vehicle, *p<0*.*0001*) antigen is significantly higher in the liver than AgV2 antigens (mean P2 = 178.5±3.09; A2 = 159.4±3.92 vs vehicle, *p<0*.*0001*) and depending on the inoculation site, the kinetics of AgV2 may differ (P2 vs A2 = 19.09, *p=0*.*0007*) (Fig 8A-B).

**Fig 7.**
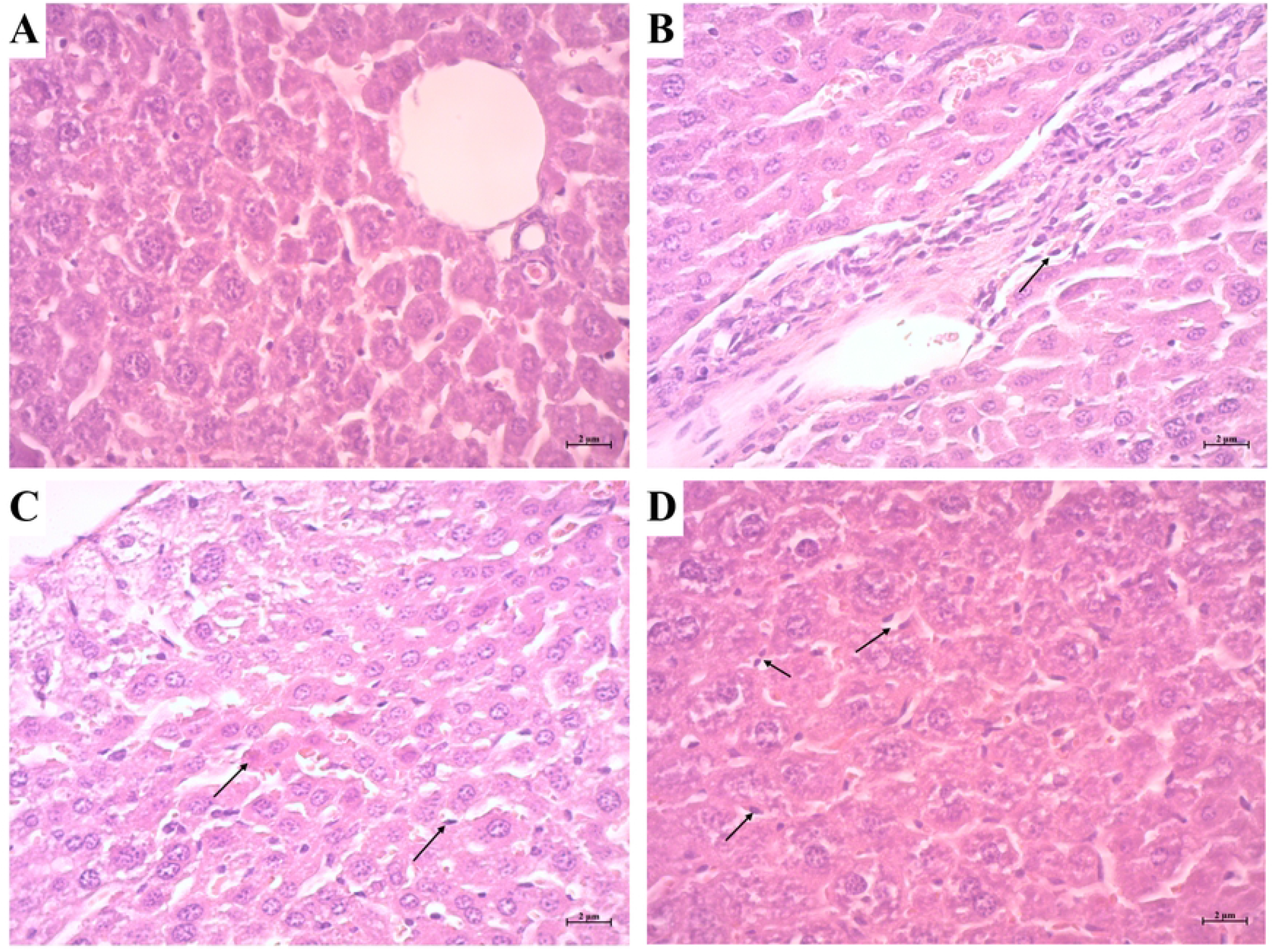
Cellular damage caused by RABV in the liver (arrows), analyzed by HE staining. A) Liver negative control. B) Infiltrative cells. C) Pyknotic cell (left arrow) and Kupffer hypertrophy (right arrow). D) Kupffer hyperplasia and hypertrophy.

**Fig 8.**
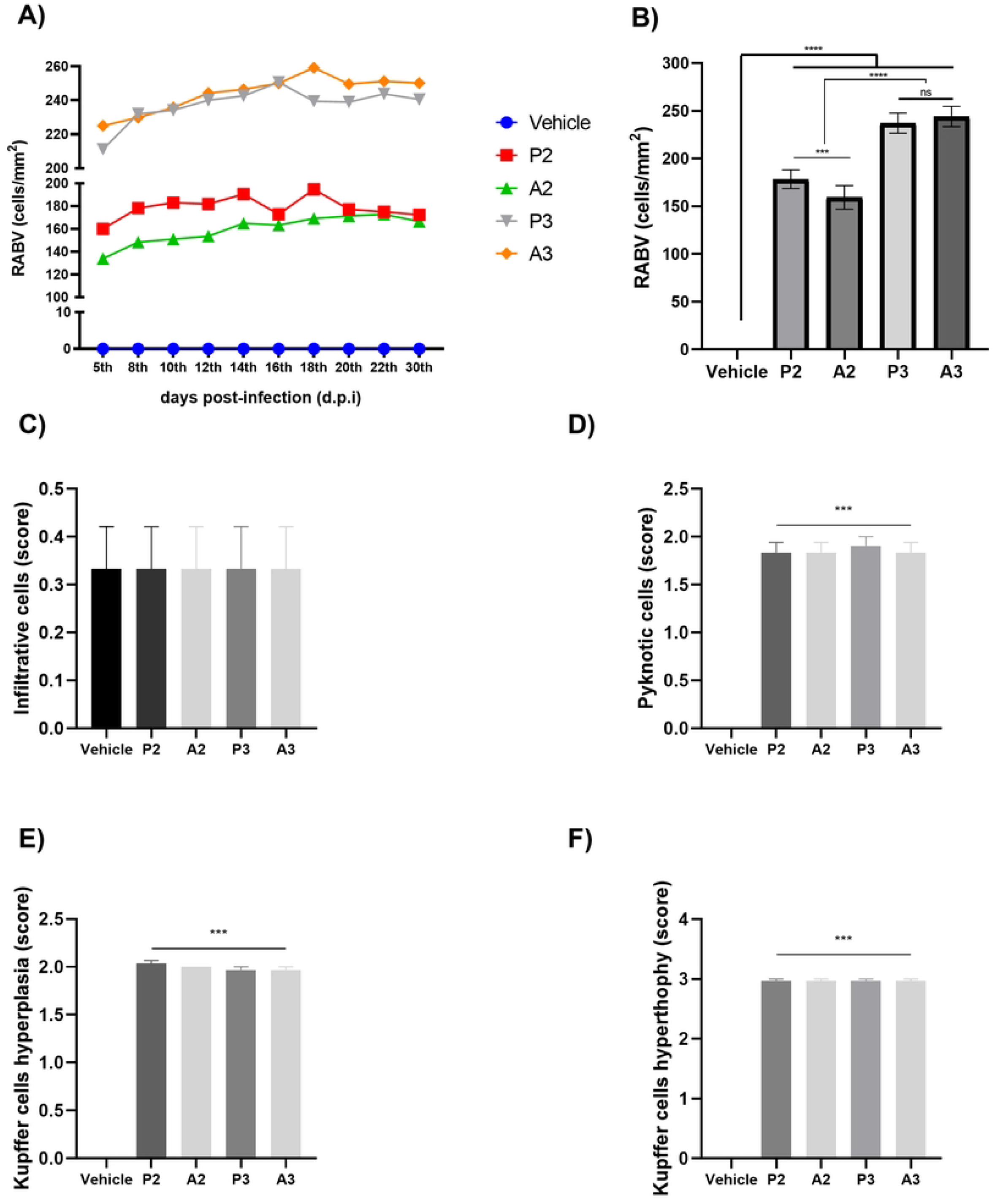
*In situ* rabies expression and RABV-induced damage in the murine liver. A) Temporal kinetics of RABV-expressing cells; B) total expression of the viral antigen; C) infiltrative immune cells in the parenchyma; D) Pyknotic cells; E) Kupffer cells hyperplasia; F) Kupffer cells hypertrophy. \*\*\**p ≤ 0*.*001*.

Contrastingly to the lungs, the infection with both strains did not change the basal scores of infiltrative cells (Fig 4C), although it was accompanied by tissue and cellular damage – with evidence of cellular death by the appearance of pyknotic morphology in the tissue (Fig 8D).

Kupffer’s cells were widely affected by both strains’ infection, as shown in Fig 8E-F since we found severe signs of hypertrophy followed by moderate hyperplasia morphology. In summary, RABV may cause tissue damage and cell death in the liver – affecting especially Kupffer’s cells.

## DISCUSSION

Although RABV is widely known as the causative agent of rabies, its infection and symptoms are not restricted to the CNS (18). However, little is known about the involvement of extraneural tissue invasion in the early and acute phases of infection and how different wild-type strains interact with other organs. Besides, we were also interested in assessing whether the immunopathological aspects were tissue-specific. Therefore, we analyzed three extraneural tissues – kidney, lung and liver – to trace the viral kinetics of different strains to infer their affinity and pathogenicity in those tissues.

Our immunohistochemistry staining demonstrated that the strains differ in their affinity to the organs, and the bat-isolated RABV strain was more expressed in all organs compared to the *wt* dog-RABV. Moreover, we could detect high levels of viral antigen - as early as five days of infection - in the lungs, liver, as previously shown (19), and especially in the kidneys – either inoculated anteriorly or posteriorly. Those results may support our hypothesis that RABV spreads to the CNS and extraneural organs simultaneously, and not necessarily retrogradely from the brain to outside the nervous system – as widely accepted. Since RABV infects the dorsal root ganglia (20) in its pathway to the CNS, we suggest the root ganglia and nerve plexuses may play a crucial role in viral dispersion within extraneural tissues.

In experimental models using the Challenge Standard Virus (CVS), which is a model of pathogenicity, the disease’s kinetics develop from the third to ninth-day post-infection – when the majority of infected mice come to death (21,22), but regarding the natural scenario, wild strains may behave differently and the incubation period varies from days to months depending on the gravity of the lesion and the incubation site. In the brain, it is also important to consider that the strain’s origin may also determine the cellular and tissue tropisms, as we showed, which might affect the course of infection and the non-neural signs.

The wide distribution *in situ* of the RABV antigen, especially in the kidney and lung, suggested that the virus might be active since the primary days (23). Despite our data did not show how the systems respond to the viral infection in the initial phase (1-4 d.p.i), which limited further analysis and conclusions, it would be interesting to trace RABV distribution by using technologies such as intravital microscopy (IVM) and confocal laser scanning microscopy – besides quantifying the viral activity by qPCR throughout the days.

Our histopathological analysis revealed an important involvement of RABV infection in the kidney, straightening the results previously published by Daher *et al*. (19), which also described kidney injury, including mild to moderate glomerular congestion, mild to severe peritubular congestion, capillary congestion, and cells in pyknosis. We could not correlate the histopathological findings to the antigen expression, as the lesions found are non-specific, however, they may be caused by hydrophobia and dehydration evoked by motor impairment of the disease. Besides, a future characterization of the cytokine and chemokine environment by using flow cytometry or immunohistochemistry would be crucial to explain the immune response and the *in situ* interactions in the renal parenchyma, which together, might integratively describe the immunopathology of RABV in the kidney.

Moreover, the renal impairment caused by RABV implies that the urine is a possible source of infection, as suggested by Debbie and Trimarchi (20) which showed the presence of RABV within the tubular cells of the kidneys. The authors also reported that mice inoculated with the urine of a fox infected with RABV died from the disease, indicating viral viability. However, the importance of urine excretions as a source of infection must be closely evaluated in a natural environment, since RABV may be inactivated by unfavorable environmental conditions.

Although rare, some studies have already described respiratory symptoms during the course of the disease in humans, which was associated with a strong activation of inducible nitric oxide synthase (iNOS) and diffusible parenchymal damage (12,24). Though the authors had failed to detect RABV genome, they suggested a close association between acute respiratory distress syndrome (ARDS) with human rabies. Contrastingly, we were able to detect the local expression of both antigenic variants’ antigens – confirming the ability of RABV to infect lung cells.

In fact, the ARDS has a multifactorial and heterogeneous aspect evoked by the pulmonary damage, which later results in a noncardiogenic edema of greater mortality (25). This damage follows the activation of resident macrophages by damage-associated molecular patterns (DAMPs) or pathogens-associated molecular patterns (PAMPs), which trigger the expression and release of inflammatory molecules and immune mediators that lead to the infiltration of adaptive and innate immune cells (26).

Moreover, the pulmonary immune response embraces the active role of orchestrated and diverse adaptive and innate immune cells which determine the outcome during the inflammatory events that affect the lungs (27). It is also essential to understand how these resident immune cells interact with the rabies virus since, in the periphery, the virus modulates the cholinergic anti-inflammatory pathway in monocytes-derived macrophages which leads to the expression of a Th2-like phenotype (28).

Here, we also indicated the moderate edema and the thickening of the alveolar wall, followed by the infiltration of immune cells, which together may suggest that this complication may participate in rabies pathogenesis in the lung of infected mice. However, the subjectivity of the symptoms and the limitations of the clinical analysis in experimental murine models limit further conclusions – but a future and more detailed characterization of the immune phenotype and the quantification of the cytokines and chemokines would reveal the involvement of the pulmonary damage in extraneural rabies. It would also be interesting to use special histological techniques to characterize profibrotic markers (29).

Regarding the histopathological findings in the liver, Kupffer’s cells were particularly affected by the infection, since we described intense hypertrophy and moderate hyperplasia, possibly indicating changes in the morphology of these cells upon RABV infection. Kupffer’s cells are resident phagocytes that are engaged in triggering the innate response in the liver sinusoid upon the interaction with pattern recognition receptors to initiate a cascade of antiviral response (30).

We argue that Kuppfer’s cells activation may have led to a pro-inflammatory response, resulting in signs of pyknosis in the liver. In spite of it, the absence of infiltrative cells may suggest that the virus limits a systemic immune reaction, which is insufficient to clear the infected cells. It is interesting because the liver displays an essential role in the maximization of anti-pathogen response by the production of acute phase proteins, and its particular microenvironment favors the tolerance of xenobiotics (31–33). Thus, assessing the production of cytokines besides characterizing the mechanisms of pathogenicity would help to understand this process.

Overall, in this study, we demonstrated that different strains of RABV have the ability to infect different extraneural organs, leading to cellular and tissue damage at different levels. We also showed the tropisms of dog and bat-variants to these organs and suggest that the kinetics of infection may develop simultaneously within the neuroinvasion, possibly through the peripheral nervous system. This scenario opens up new perspectives regarding rabies pathogenesis and could contribute to the development of new therapeutic approaches.

## MATERIALS AND METHODS

### Ethics

The Internal Ethics Committee on the Use of Animals for Research of Evandro Chagas Institute (CEUA/IEC) approved this project under the license no. 0011/2009. Experimental procedures were carried out under the guidelines of the Brazilian Council for Control of Animal Experimentation (CONCEA).

### Viral Stock Production and Titration

To produce the viral stock, two RABV-positive samples were selected: a canine-strain antigenic variant 2 (AgV2) and a bat-strain antigenic variant 3 (AgV3). Both samples were stored in an ultrafreezer at Rabies Laboratory of the Evandro Chagas Institute, Brazil. These samples were diluted in penicillin and streptomycin (Pen-Strep) (1:2) and salin (1:1); and then intracerebrally inoculated in newborn swiss mice. After the start of paralysis, the brains of moribund animals were collected and proceeded to dilution in albumin-bovin solution containing antibiotics to determine viral titers in the sample. Based on the Reed & Muench method (34), the AgV2 titer determined was 4.9 LD50/0.02mL and the AgV3 titer was 5.4 LD50/0.02mL.

### Experimental Infection

A total of 155 Swiss albino mice (*Mus musculus*) were categorized into five experimental groups. Intramuscular injections were administered in either the posterior or anterior region of the animals. A total of 70 mice were inoculated with antigenic variant 2 (AgV2), while 70 mice were inoculated with AgV3, and 15 mice were administered with the vehicle solution (albumin-bovin solution containing antibiotics). Kidneys, lungs, and livers, were collected from the animals on days 5, 8, 10, 12, 14, 16, 18, 20, 22 and 30 post-infection (d.p.i), subsequent to the inoculation process (fig. 9).

**Fig 9.**
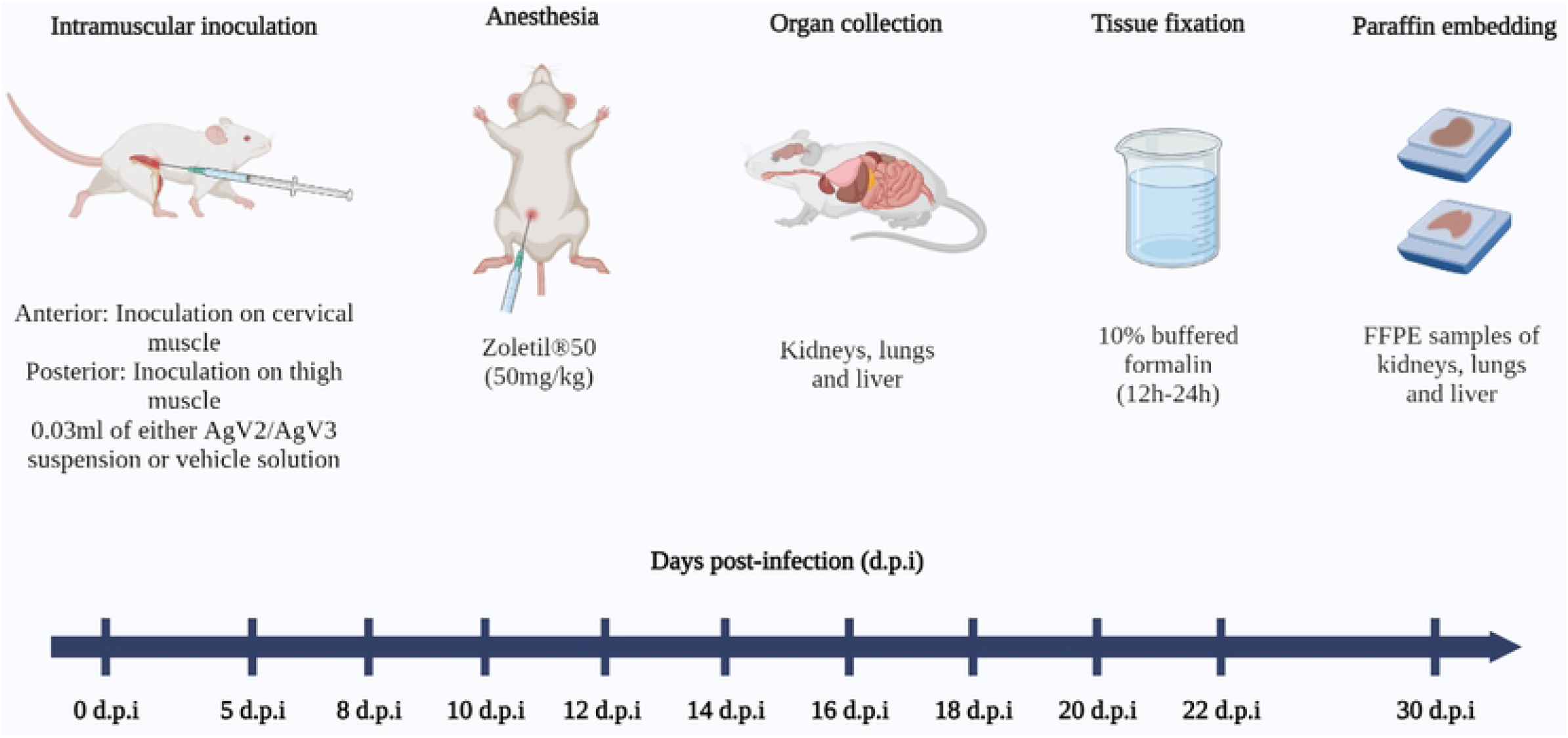
Experimental design and organ collection. The figure illustrates each procedure performed, from viral inoculation to sample collection and tissue preparation throughout the kinetics (0-30 days post-infection).

### Histopathological Processing

Collected organs were fixed in 10% buffered formalin solution for 12 to 24 hours, dehydrated in graded ethanol series (70%, 80%, 95%, and 100%), followed by twice-purification with xylene, and then twice submerged in paraffin at 60 ºC. After the histopathological processing, we embedded the organs in paraffin. The tissues were sectioned into 5µm sections, which were later stained with hematoxylin-eosin and analyzed with a light microscopy, according to the routine of the Laboratory of Pathological Anatomy of Evandro Chagas Institute.

### Histopathological Analysis

We conducted the histopathological analysis considering cellular, structural and circulatory alterations in the kidney, lung, and liver. We performed a semi-quantitative analysis based on the degree of the lesion. A score ranging from 0 to 3 was assigned, with 0 being considered a normal histology pattern or absence of the lesion; 1 for mild injury; 2, for moderate injury; and 3 for severe injury.

### Immunohistochemistry

We deparaffinized the tissues in xylene for 25 minutes, followed by tissue hydration in decreasing concentrations of ethanol: 100%, 95%, 80%, and 70% for a total time of 14 minutes. Blocking of endogenous peroxidase was performed in 3% hydrogen peroxide for 45 minutes. We performed antigen retrieval with citrate buffer (pH 6) + Tween®20 in a water bath at 95 ºC for 20 minutes. Nonspecific proteins were blocked using 10% non-fat milk for 30 minutes at room temperature. Finally, the slides were incubated overnight at 4 ºC with the anti-RABV primary antibody (1:50 dilution, produced in-house) diluted in 1% bovine serum albumin (BSA). Tissues were then incubated with biotin and streptavidin for 30 minutes each at 37 ºC. Antigen-antibody reactions were revealed by diaminobenzidine (DAB) chromogen. Finally, the tissues were counterstained with Harris hematoxylin, followed by dehydration with increasing concentrations of ethanol (70%, 80%, 95%, and 100%) and xylene. Lastly, the slides were dried off and mounted.

### Statistics

The immunohistochemical staining pattern was analyzed by quantifying all positive cells in 10 fields in a light microscope. The results were analyzed by calculating the means and the standard error; and then statistically analyzed by the one-way and two-way ANOVA test, followed by the Bonferroni test, assuming statistical significance for p<0.05. Statistical analyzes were processed on GraphPadPrism version 8.0 for Windows (GraphPad software, San Diego, CA, USA).

## ACKNOWLEDGEMENTS

We would like to thank Orlando Pereira and Valter Campos, from the Department of Clinical and Experimental Pathology (SAPEX) of Evandro Chagas Institute (IEC), for providing technical support.

